# Coding of egocentric distance in the macaque ventral intraparietal area

**DOI:** 10.1101/2024.07.31.605976

**Authors:** Baptiste Caziot, Sadra Fathkhani, Frank Bremmer

## Abstract

The encoding of three-dimensional visual spatial information is of ultimate importance in everyday life, in particular for successful navigation toward targets or threat avoidance. Eye-movements challenge this spatial encoding: 2-3 times per second, they shift the image of the outside world across the retina. The macaque ventral intraparietal area (VIP) stands out from other areas of the dorsal ‘where’ pathway of the primate visual cortical system: many neurons encode visual information irrespective of horizontal and vertical eye position. But does this gaze invariance of spatial encoding at the single neuron level also apply to egocentric distance? Here, concurrent with recordings from area VIP, monkeys fixated a central target at one of three distances (vergence), while a visual stimulus was shown at one of seven distances (disparity). Most neurons’ activity was modulated independently by both disparity and eye vergence, demonstrating a different type of invariance than for visual directions. By using population activity, we were able to decode egocentric distance of a stimulus which demonstrates that egocentric distances are nonetheless represented within the neuronal population. Our results provide further strong evidence for a role of area VIP in 3D space encoding.

## Introduction

The Posterior Parietal Cortex (PPC) is composed of many areas that are anatomically and physiologically distinct^1^. Neurons respond to a variety of signals that are largely redundant across PPC areas, and their respective functional roles remain elusive. Clinical observation has consistently showed that lesions to the PPC strongly impact the generation and guidance of spatial awareness^2^. Moreover, these deficits are especially severe in the depth dimension (e.g. ^3–5^ for earlier studies, but see also ^6^ for recent evidence in that direction), but this link has never been well established at a physiological level. The Ventral Intraparietal area (VIP), located in the fundus of the intraparietal sulcus, stands out from other areas of the dorsal pathway in one respect: many neurons in this area encode spatial information invariant of horizontal and vertical eye position^7–9^. Such invariance at the single cell level is different from population-based approaches which have been suggested for ensembles of neurons carrying an eye-position signal (gain field)^10–13^.

A similar invariance problem exists in the depth dimension: ever since the invention of the stereoscope by Wheatstone^14^, it has been well established that the difference in projection of a single point on the two retinae (binocular disparities) is the source of the clear sense of depth that one generally experiences when looking at the environment (stereopsis)^15–18^. Geometrically, binocular disparities depend on the relative distance between a point and the current angle formed by the eyes (vergence). Therefore, binocular disparities alone are insufficient to sustain the perception of the absolute distance of a point to the observer, or egocentric distance. While today there is good agreement that observers are able to perceive egocentric distances from binocular vision, including at far distances^19–22^, where and how this information is computed in the brain remains unknown.

Theoretically, there are at least two ways that egocentric distances could be encoded in the brain: either as an explicit encoding through a mechanism equivalent to disparity scaling^15^, or implicitly through a distributed population code^23^. Some studies have found modulations of neural activity by vergence in early visual areas^24,25^, although these results have not been consistently replicated^26,27^. Furthermore these modulations were highly non-linear, making it unclear how they could be combined at a population level^15^. In humans, disparity tuning has been observed in various visual cortical areas including in the PPC^28–30^. Likewise, in macaque monkeys, disparity tuning has been showed in numerous visual cortical areas, starting in the primary visual area V1 and reaching up to the highest levels of the dorsal stream: the Lateral Intraparietal area ^31,32^, Ventral Intraparietal area (VIP)^33,34^ and extending into sensorimotor systems including the Parietal Reach Region^35^, the Anterior Intraparietal area^36,37^ and area V6^38^. In contrast, while *conjugate* eye position signals are omnipresent in the PPC^11,39,40^, *disconjugate* (vergence) eye-position signals have rarely been investigated. Although some studies did report modulation of neural activity by vergence^31,32,35,41^, suggesting that the PPC could, in principle, encode egocentric distances.

Since area VIP demonstrates invariance of spatial locations to conjugate (2D) eye-positions, here we ask whether this invariance extends to disconjugate eye-positions (depth). We investigated neural activity in area VIP in response to stationary and moving stimuli at various egocentric distances, while animals fixated at different distances. We found that neural activity is modulated by stimulus motion direction, stimulus disparity and vergence angle (fixation distance). To further characterize single-cell activity, we modelled neural activity using a Linear-Nonlinear Poisson model^42,43^. By inverting this model, we were able to recover egocentric distances of our stimuli, with biases consistent with the psychophysical literature. Our results suggest that egocentric distances are encoded implicitly through a heterogenous population code and provide strong evidence for a role of area VIP in 3D space encoding.

## Results

### Task

Two macaque monkeys (*macaca mulatta*) were trained to keep fixation on a central target. The target had one of 3 possible egocentric distances corresponding to vergence angles of 1.5, 3.5 and 5.5 deg (see Figure 1A, B). After an 800 ms fixation period, a static Random Dot Pattern (RDP) was presented dichoptically for 250 ms, after which it started to move at a constant speed for 1 and 1/4 cycle of a circular path in the fronto-parallel plane (see Figure 1C). Below we refer to these three temporal intervals as the *fixation period, stationary period* and the *motion period*. The RDP had one of 7 egocentric distances (ranging from 27 to 229 cm in front of the animals), resulting in multiple combinations (each termed a *condition* in the following) of disparities relative to fixation ranging from -5 to +5 deg, in steps of 1 deg, with a negative sign corresponding to crossed disparities (closer than fixation). Eye-positions were recorded using scleral search coils and animals successfully maintained fixation throughout the duration of the trials for liquid reward (see Figure S1). We recorded neural activity from 113 individual neurons (89 and 24 for animals C and H respectively), with an average number of 9±2 trials per cell and condition. In the following, we analyze first neural tuning for basic features (motion direction, disparity and vergence), before turning to egocentric distance encoding.

**Figure 1:**
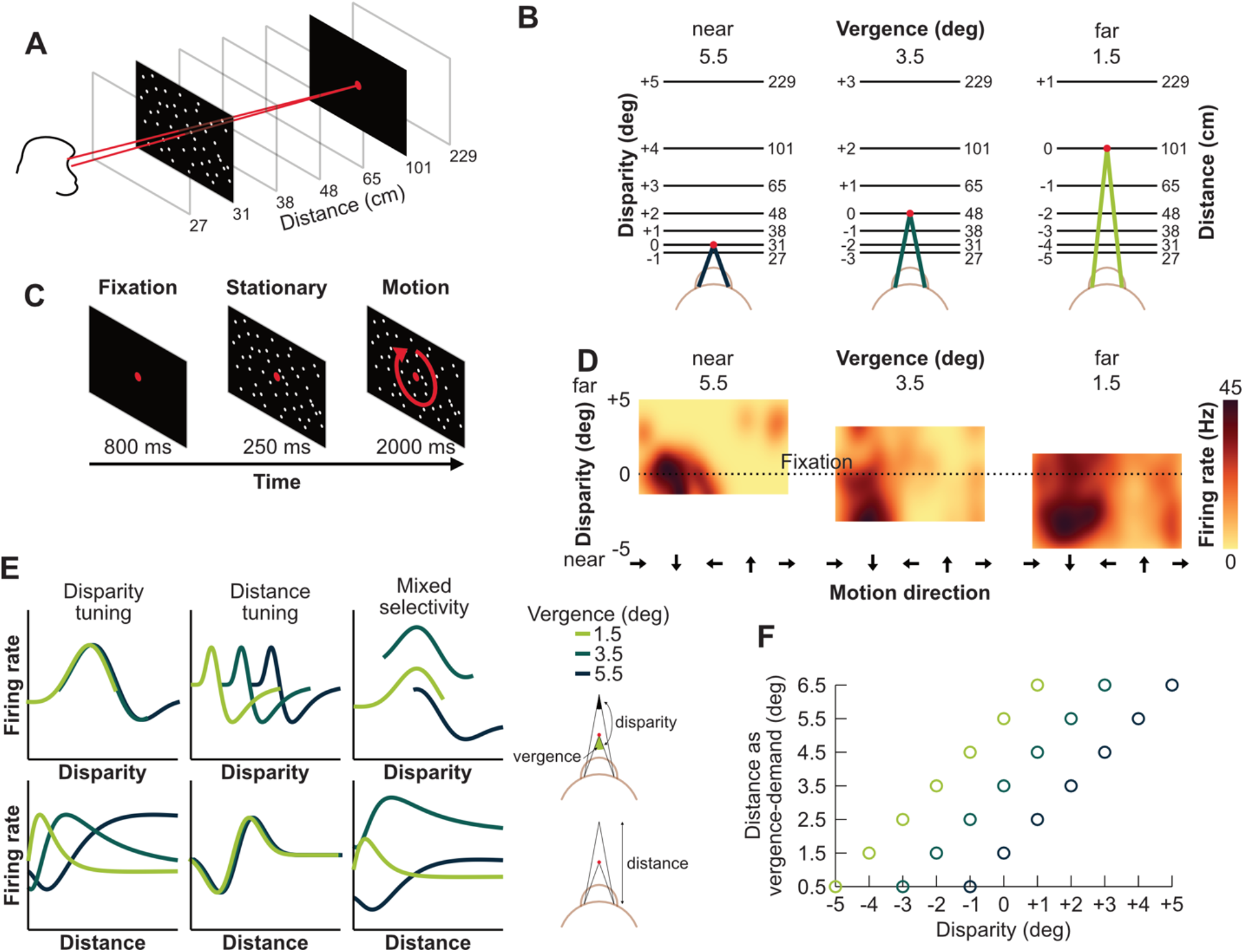
**A)** Monkeys were trained to fixate a central target while a Random Dot Pattern (RDP) was shown stationary or moving in the fronto-parallel plane at one of seven possible distances (shown in centimeters at the bottom). **B)** The fixation target (red dot) could have one of three possible distances to the monkeys, requiring different vergence angles (shades of green). Horizontal lines represent the possible stimulus distances. Numbers on the left are disparities in deg and on the right egocentric distances in cm. Note that the stimulus distances are always the same, but their disparity depends on fixation distance. **C)** Time-course of a trial. Initially, only the fixation target was shown for 800 ms. Then, a static RDP was displayed for 250 ms, after which it started moving at a constant speed on a circular pathway^44^ for 2000 ms, completing 1 and 1/4 cycle. **D)** Peristimulus Time Histogram of an example neuron as a function of motion direction (abscissa), disparity (ordinate) and vergence angle (panels). Colors represent firing rate in Hz. **E)** Hypothetical neuronal tuning curves. Each column represents one type of encoding as a function of disparity (top) or egocentric distance (bottom). The neuron in the first column would be tuned to disparity irrespective of egocentric distance. The neuron in the second column would be tuned to egocentric distance irrespective of disparity. And the neuron in the last column would exhibit mixed selectivity to both disparity and vergence. Figures on the right depict the relationship between vergence, disparity and egocentric distance. **F)** Representation of the stimuli geometry used in this experiment. Here distance is represented as vergence-demand (binocular parallax), that is the vergence angle that would be required to fixate at this distance. In this space, a simple relationship exists between vergence, disparity and vergence-demand: vergence demand = vergence - disparity.

### Direction tuning

A dominant feature of area VIP is a high degree of selectivity for motion direction^44,45^. All neurons demonstrated significant directional tuning for fronto-parallel visual motion in at least one of the 21 (3 vergence x 7 disparities) motion period conditions (p<0.05, uncorrected for multiple comparison). To estimate the consistency of directional tuning across disparity^33,46^ and vergence, we computed pairwise differences of preferred directions across the 21 conditions for each cell. Figure 2B (bottom, “PD differences”) plots the distribution of these differences. This distribution was more narrowly centered on 0 than the same distribution computed on a randomly shuffled dataset (Figure 2B, bottom, “Shuffled”), and their medians were significantly different indicating that preferred directions are more stable across conditions than expected by chance. We also compared this distribution to what should be expected from measurement noise by resampling the dataset within conditions (Figure 2B, bottom, “Measurement noise”), that is for identical visual stimuli where we expect preferred directions to remain perfectly stable. The median of the measured distribution was not significantly different from the median of the measurement noise distribution, demonstrating that neurons’ preferred motion direction was constant across vergence and disparity.

**Figure 2:**
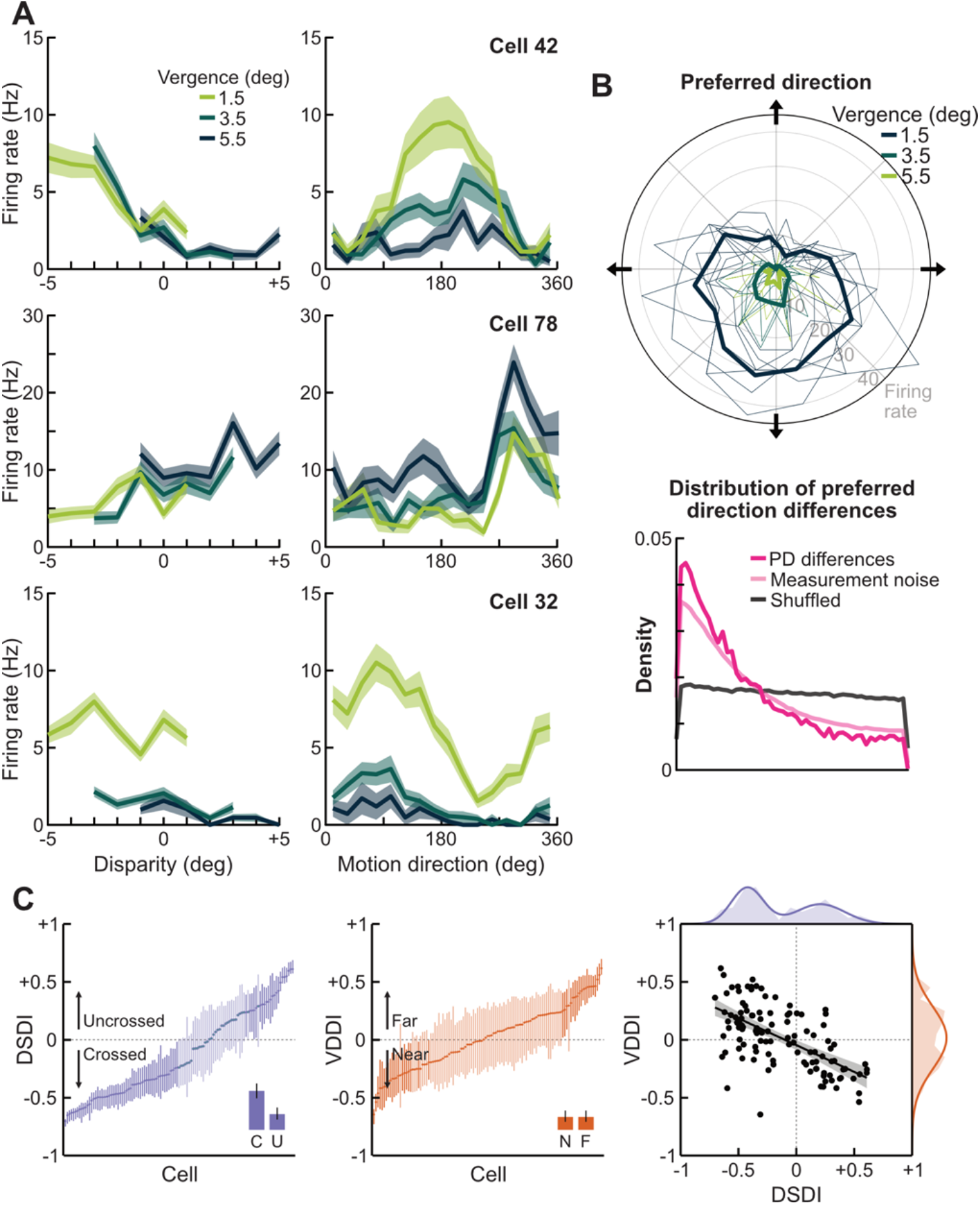
**A)** Examples of neurons’ tuning curves in the motion period (excluding the first ¼ of cycle). Each row displays the activity of an individual neuron. Left column shows activity as a function of disparity marginalized over directions. The right column shows activity as a function of direction marginalized over disparity. Colors represent different fixation distances. Cells 42 and 78 show a preference for near and far disparities respectively, and their activity was largely unaffected by vergence angle. In sharp contrast, the activity of cell 32 is clearly modulated by vergence angle. **B)** Top: Direction tuning of a neuron. Colors represent different fixation distances. Thin lines are activity for each distance condition separately and thick lines are averages. Bottom: Distribution of differences in Preferred Direction across conditions for all the neurons recorded from both animals (pink). The black distribution shows differences in preferred directions in a randomly shuffled dataset, and the light pink distribution shows differences in preferred directions when the dataset is resampled within conditions (for identical visual stimuli), indicating expected measurement noise when preferred directions are perfectly stable. **C)** Left: Depth Sign Discriminant Index (DSDI) for all cells sorted by magnitude. Saturated lines indicate neurons whose 95% confidence did not include 0. A negative DSDI indicates a preference for crossed disparities (near), and positive for uncrossed disparities (far). Insets represent the fraction of neurons with a significant preference for crossed (C) vs uncrossed (U) disparities, and 95% Confidence Intervals. Middle: Vergence Distance Discriminant Index (VDDI). Graphical details are identical to the left subfigure. Here a negative VDDI indicates a neuron that responds more during near fixations than far fixations, and conversely for a positive sign. Right: VDDI plotted as a function of DSDI for each neuron (black dots). Black line shows regression line and shaded area confidence interval. On the top and right are marginal distributions (shaded areas) and best Gaussian Mixture Model fit (thick lines).

### Disparity tuning

Another characteristic feature of area VIP is a clear selectivity for binocular disparities, including a preference for near (crossed) disparities^33,34^. In this dataset, 55% of the neurons showed a significant tuning for disparity during the motion period (Kruskal-Wallis test, p<0.05, uncorrected for multiple comparisons). We assessed their preference for crossed vs. uncrossed disparities by computing a Depth Sign Discriminant Index (DSDI)^47,48^. A DSDI of -1 indicates a preference for crossed disparities, of +1 a preference for uncrossed disparities and of 0 a cell that responds equally to crossed and uncrossed disparities. This DSDI index was significantly different than 0 for 51% of the neurons (permutation test, p<0.05, uncorrected for multiple comparisons). Among this subpopulation of neurons, 78% (45/58) exhibited a preference for crossed (near) disparities (Figure 2C, left), in good agreement with previous results^33,34^. The distribution of DSDI values appeared bimodal (see Figure 2C, right), suggesting that it reflects a binary class of disparity receptors. We tested this hypothesis by fitting a Gaussian Mixture Model (see Methods) to the distribution of DSDIs and found that a model including 2 components fitted data better than a model that assumed a unimodal distribution of DSDIs.

### Vergence tuning

Finally, we examined neurons’ selectivity to fixation distances (vergence), a feature that remains unknown in area VIP. Since in our dataset the range of simulated fixation and stimulus distances was fixed, vergence and disparities are unavoidably anti-correlated (see Figure 1F). Consequently, we only included comparable disparities (−1, 0 and +1 deg relative to fixation distance) in these analyses. We found that 33% (37/113) of cells were significantly tuned to vergence during the motion period, 17% (19/113) during the stationary period, and 26% (29/113) during the fixation period where only the fixation target was displayed (Kruskal-Wallis test, p<0.05, uncorrected for multiple comparisons). Out of the entire dataset, 11 cells remained significantly tuned for vergence throughout the 3 stimulus periods. Therefore, while selectivity for fixation distance is not as pronounced as motion or disparity in area VIP, it remained well above chance level.

For each cell we then computed a Vergence Distance Discriminant Index (VDDI, see Methods), by analogy with the Depth Sign Discriminant Index. This index varies between -1, corresponding to a preference for near fixation distances (within the range of distances tested within this experiment), and +1 to a preference for fixation at far distances; here again restricting analyses to matching disparities across vergence angles. 28% (32/113) of cells had a VDDI significantly different than 0 (see Figure 2C, middle; permutation test, p<0.05). In contrast with disparity, there was no clear asymmetry between near and far fixation distances, and the distribution of VDDI values appeared unimodal (see Figure 2C, right). A Gaussian Mixture Model that includes only 1 component fitted data better than a model assuming a bimodal distribution (see Methods). Finally, there was a clear and significant anti-correlation (resampling, ρ = -0.62, p<0.001) between the cells’ disparity preference (DSDI) and cells’ fixation preference (VDDI), even though we only included conditions with identical ranges of disparities in this analysis. Put differently, cells selective for fixation at closer distances tend to be selective for stimuli with uncrossed disparity (farther than fixation), and conversely for cells selective for fixation at farther distances.

### Egocentric distance

As a next step, we aimed to determine if neural responses to stimulus disparity and vergence were independent or not at a single cell level. If neurons are tuned to absolute distance, they should exhibit stable activity when stimulus distance remains constant (see Figure 1E, middle column), despite changing disparities and vergence. In contrast with this hypothesis, we found that 69% of the neurons (77/113) significantly changed their activity with constant egocentric distance (Kruskal-Wallis test, p<0.05, uncorrected for multiple comparisons). Among the remaining 36 neurons, only 4 demonstrated significant response to a specific distance (Kruskal-Wallis test, p<0.05), and none when applying a Holm-Bonferroni correction for multiple comparison.

The opposite reasoning is that if a neuron is tuned to the egocentric distance of a stimulus, its tuning for disparity should invert as fixation distance crosses the neuron’s preferred egocentric distance. Although we did not test all possible distances within the visual scene, our stimuli included a broad range of egocentric distances and such inversion should be evident in at least some cells. We resampled our dataset for each neuron separately and computed the DSDI for the smallest and largest vergence angle on each sample. We then computed the 95% two-sided confidence interval of the difference of DSDI between the 2 vergence. This confidence interval did not include 0 for less than 4% of the neurons (4/113), and none of the neurons when applying a Holm-Bonferroni correction for multiple comparison.

In summary, we found no evidence for a direct representation of egocentric distance by individual neurons in area VIP (shifting of disparity tuning curves by vergence angle): almost all neurons varied their activity when disparity or vergence changed but the stimuli maintained their egocentric distance, and no neuron inverted its disparity sign preference with vergence angle. Yet, we still found a tuning for both disparity and vergence angles, which suggests that egocentric distance is encoded at a population level. In the next sections, we investigate how reliably these variables, including egocentric distance, can be decoded from population activity.

### Encoding model

To better characterize the neuronal tunings, we fitted a Linear-Nonlinear Poisson (LNP) spiking model to our data^42,43^. This class of models has previously been showed to account well for activity in the PPC^42,49^. This model assumes that task variables contribute additively to the neurons firing rate then passed through a static nonlinearity (see Figure 3A). While the source and shape of this static non-linearity remains unclear, differences in firing rate of individual neurons when two variables co-vary is linearly related to the differences in firing rate when each variable varies individually on a logarithmic scale (see Figure S5), demonstrating that an exponential non-linearity approximates neurons’ activity well.

**Figure 3:**
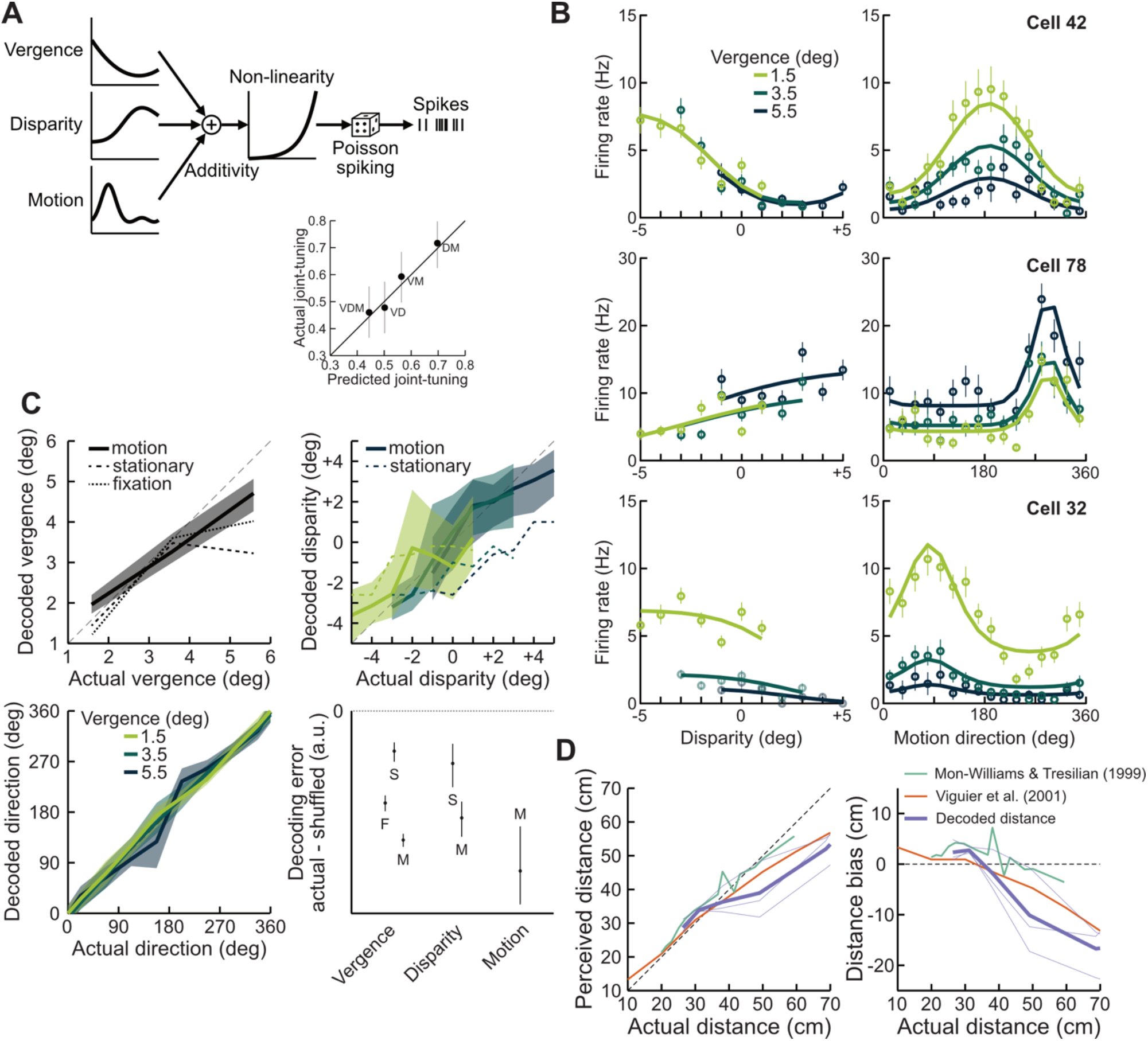
**A)** Depiction of the Linear-Nonlinear Poisson model. Variables (Vergence, Disparity and Motion) contribute additively to the activity of the neuron and are then passed through a static exponential nonlinearity. Bottom inset shows fraction of neurons tuned jointly to combinations of variables (for example DM corresponds to a joint tuning to Disparity and Motion), as a function of fraction of joint-tuning predicted from an independent tuning to each variable. **B)** Example of predicted tuning curves for the same 3 neurons as in Figure 2. Solid lines are best model fit to the neural responses (+/-SEM) as shown by individual points. **C)** Decoded vergence (top-left), disparity (top-right) and motion direction (bottom-left) during the Motion period as a function of the stimulus values. Colors show different fixation distances and shaded areas are 95% confidence interval of decoded values. Dashed lines show median decoded values during the Fixation and Stationary periods (see Figure 1C), which were not fitted to our model. Bottom-right subplot shows median and 95% Confidence Interval difference decoding error (distance to true value) between the actual and randomly shuffled dataset, for all 3 variables and different stimulus periods (F=Fixation, S=Stationary, and M=Motion). A negative number indicates a better decoding accuracy than expected from chance. Note that units for different variables are different and cannot be directly compared. **D)** Egocentric distance prediction during the Motion period (purple) as a function of stimulus distance (abscissa) and comparison with results from human data from Mon-Williams & Tresilian (1999, orange)^19^ and Viguier et al. (2001, green)^20^.

To assess which variables contribute to the activity of each neuron, we used a cross-validated forward nested-search approach (see Methods). In short, each variable (motion, disparity, vergence) was added sequentially. The variable that most improved the likelihood of the model, estimated on an unfitted test-set, was preserved. We repeated this process until no more variables improved the likelihood of the model. This approach let us estimate jointly which variables contribute to the neuron’s activity and the shape of their tuning curves. Figure 3B shows examples of model fits for the same neurons as in Figure 2.

Results are in good agreement with previous analyses. With this approach we found that 89% of cells are tuned to motion, 81% to disparity and finally 64% to vergence. Predictions from model fits were able to replicate all results from previous analyses (see Figure S6), demonstrating that it provides a good description of a neuron’s activity. The overall correlation between actual and predicted firing rates was 0.91. The correlation between the DSDI and VDDI measured and predicted from the model were respectively 0.99 and 0.94. We found no evidence for a joint coding of variables above chance level (see Figure 3A), suggesting that the encoding of these variables is homogeneously distributed throughout the population.

### Decoding

The LNP encoding model generates probability functions of neural activity based on task variables, and can be inverted to predict task variables given the population activity. Specifically, for each neuron, we can compute the probability of each combination of task variables given the neurons’ current firing rate. We can then determine the stimulus values that are most likely to have produced the neural activity across the population. Since cells were recorded separately, we simulated population activity by randomly picking 1 trial from identical disparity and vergence conditions independently for each cell 10,000 times and decoded the most likely stimulus values. Figure 3C shows decoding results for all 3 task variables (motion direction, disparity and vergence) during the motion period.

We could decode vergence angle with a mean precision of 0.15 degree (SD of the distribution of decoded values). Decoded vergence was approximately unbiased for the fixation distance of 3.5 deg, but was significantly overestimated for a vergence angle of 1.5 deg (far fixation) and underestimated for a vergence angle of 5.5 deg (near fixation; median decoded vergences respectively 1.96 and 4.71, p<0.001).

We found that we could similarly decode disparity with a good level of precision. The mean error was 0.67 deg. Decoded disparities were significantly underestimated at the most extreme disparity values (−5 and +5 deg, decoded values respectively -3.6 and +3.5, p<0.001), as well as for the mean bias over crossed and uncrossed disparities (mean bias respectively +0.45 and -1.0). However, there was no statistical difference between the mean errors for crossed vs. uncrossed disparities (resampling of the mean error, p=0.85). We also found no difference between decoding precisions at the different fixation distances.

Decoded motion direction was excellent: the mean precision for motion direction was 8 deg and we found no interactions with vergence or disparities.

Our model was fitted to neural data during the motion period exclusively. However, if these variables are encoded by the population of neurons and their tuning curves remain relatively stable across stimulus periods, we should be able to decode the stimulus variables during the other, unfitted, stimulus periods. These results are plotted alongside data from the Motion period in Figure 3C. Decoding performance (both precision and accuracy) were unsurprisingly significantly poorer than during the motion period. Yet, we compared decoding accuracy (difference to true variable value) with that of a randomly shuffled dataset and found that we could decode task variables above chance level for all 3 stimulus periods (see Figure 3C, bottom-right).

In summary, our results show that information about motion direction, stimulus disparity and vergence can be decoded from population activity in VIP. We found that decoded vergence was approximately unbiased at a distance of 48 cm, but were overestimated for near fixations (farther than fixation) and underestimated at far fixation (closer than fixation). Similarly, decoded disparities were consistently smaller than their true value (closer to fixation). We used these variables to recover egocentric distances of the stimuli. Figure 3D shows the decoded distance during the motion period. Decoded distances were overestimated for the nearest stimuli. This was true for all vergence angles for stimuli at both 27 and 31 cm. Inversely, decoded distances were underestimated for all vergence angles for any stimuli farther than 48 cm. These results should be compared to psychophysical measurements of human distance estimation. Figure 3D plots results from 2 studies that explicitly measured binocular distance perception. Notably, both studies found, as in our results, an overestimation of close distances and an underestimation of far distances, with an inversion of this bias somewhere between 40 and 50 cm. These biases are also comparable in amplitude to those measured in humans.

## Discussion

The ventral intraparietal area (VIP) stands out from other areas of the dorsal pathway of the macaque visual cortical system: many neurons encode spatial information invariant with respect to horizontal and vertical eye position^7,8^. Here we asked if such eye position invariance is also found for distance. Such an invariance would correspond to a shift of disparity-tuning curves by vergence angle (see Figure 1F). Our data do not provide evidence for this hypothesis at the scale of individual neurons. Instead, we found that individual neuron’s activity was well described by a Linear-Nonlinear Poisson model (LNP), and that a gaze invariant encoding of egocentric distance can be achieved at a population level. This finding is important for at least two major reasons: first, the role of area VIP for behavior is yet not fully understood. It has been implicated in goal-directed navigation^33,44^ (visual–vestibular interactive responses) but also threat detection and/or avoidance^50–52^. Both behavioral contexts require a robust estimation of egocentric distance. Our data clearly show that the relevant information is multiplexed in the population activity of area VIP. Second, a functional equivalent of macaque area VIP has been identified in humans^53,54^. Hence, we can infer from our current findings that the neural encoding in the posterior parietal cortex of humans also contains such information.

### Individual tuning

Almost all neurons exhibited selectivity for motion direction. Consistently with previous studies^33,34^, we found that directional tuning in area VIP was unaffected by stimulus disparity. Here, we extend this finding by showing that this directional tuning is also unaffected by changes in vergence angle (Figure 2B). This new finding further demarcates the functional roles of area VIP as compared to area MST, since cells in the latter often flip their directional selectivity with disparity^34,46,55^. Furthermore, 78% of the neurons significantly tuned to non-zero disparities preferred crossed (near) disparities. This result confirms previous findings that neurons in area VIP have a preference for near extrapersonal space^33,34,45^, regardless of fixation depth. A similar overrepresentation of crossed disparities has been found in other areas of the PPC^31,32,35^. A smaller, but consistent, overrepresentation of crossed disparities has also been observed in early visual areas. Sprague et al. (2015)^56^ have argued that this bias is a byproduct of the oversampling of cells from the lower hemifield. The same argument is unlikely to apply for the PPC, first because the overrepresentation of crossed disparities is far stronger, second because receptive fields in the PPC are very large, with very few cells being exclusively selective to the upper or lower hemifield^7,44,45,57^, and finally because area VIP is not organized in a retinotopic manner^54^ making it less likely that receptive fields from the lower visual hemifield have been oversampled.

We have found no clear preference for fixation distances across the population. However we found a negative correlation between vergence and disparity (Figure 2C), a characteristic previously reported in LIP neurons^32^. This result is not extremely surprising, considering the inverse relationship between fixation distance and disparities within the visual scene. For instance, when fixating at infinity all points within the visual scene have crossed disparities, so it would not make sense for a neuron highly selective for fixation at infinity to also be selective for uncrossed disparities. Nonetheless, this finding demonstrates that the tuning characteristic of individual neurons is not random; which might be evidence of a coding scheme, such as efficient coding.

### Eye-position signals

We have shown that neurons activity in area VIP are modulated by vergence eye-posture. While these modulations were small, their magnitude (Spikes/deg) is comparable with modulations that have been showed for version (conjunctive) eye-position signals throughout the cortex^8,11,39,58–61^, and allowed decoding vergence eye-posture with a reasonable level of precision. Furthermore, these small modulations are amplified by the non-linear output of neurons. In particular, neurons in area VIP tend to prefer higher motion speeds than used in our study^45^. We can predict that the small but consistent vergence modulations reported here would be dramatically amplified at higher motion speeds. Finally, we found that this representation was stable enough that we could decode vergence eye-posture at all stimulus periods, even though our model was fitted to neural activity during the motion period only.

Theoretically, three signals could contribute to this vergence signal: sensed eye-position^62,63^, efference copy/corollary discharge signals^11^, or vertical disparities^64,65^. All three cues have been showed to contribute to perception^66–68^ or have been suggested to be the neural basis of perceptual localization^11^. Here we used stimuli where cues were congruent, and consequently we cannot dissociate which one contributed to the eye-position signals. Furthermore, we only used symmetrical vergence eye-postures (straight-ahead). In principle, vergence could be represented as a proper vergence eye-posture signal, or as a combination of monocular eye-position signals. Further investigations, using a broader range of vergence and version eye-positions, and especially asymmetrical eye-postures, are required to answer this question.

It should be noted that this result does not directly imply that observers have conscious access to their vergence eye-posture^69^. Indeed, many visual cues are known to be used by the brain without conscious access to them. It would not even be unique in binocular vision alone, where observers have poorer access to absolute disparity information than relative disparity information^70,71^ as well as no conscious access to vertical disparity information^72^.

### Distance estimation

We did not observe distance-tuned neurons in our population (neurons that shift their tuning curves based on vergence, see Figure 1F). Yet we have shown that it is possible to recover egocentric distances at a population level, based on vergence and disparity signals. That binocular disparities alone are insufficient to recover egocentric distance has long been appreciated^73^, starting with Wheatstone himself^74^. Whether observers are able to perceive egocentric distances based on binocular vision has been the subject of a longstanding debate initiated by Helmholtz and Hering^73^. Today there is a general agreement that egocentric distances can be estimated from binocular vision (but see^69^ for a new take on Hering’s point of view). Psychophysical studies have consistently found biases in distance estimation with an overestimation of close distances, and underestimation of far distances^19–22^. We have found that these biases are consistent with our decoded results.

In any case, we have shown that both disparity and vergence signals coexist within the same area of the PPC. It would be surprising that the visual system does not make use of these signals while they are readily accessible.

### What is area VIP for?

Area VIP is not typically considered as part of the classical pathway for binocular vision^15–18^. Yet we found that in this area a majority of neurons are tuned to both disparity and vergence. Binocular disparities can be discriminated just as quickly as luminance^75,76^ and are sufficient to sustain the orienting of attention in depth^77^. Furthermore at least in some circumstances, crossed (near) disparities more potently orient attention^78,79^. Finally behavioral and physiological studies suggest a link between spatiotopic representations and the orienting of attention^80^. Since area VIP is (1) one of the areas whose activity is most strongly modulated by attention^81^; (2) has cells with spatiotopic, or at least craniotopic, receptive fields^7,8^; and (3) exhibits a strong over-representation of crossed disparities^33,34^; then it seems likely that area VIP is involved in the orienting of attention in 3-dimensional space and its role for the maintenance of 3-dimensional perceptual stability.

## Conclusion

In conclusion, we have shown that neurons’ activity in area VIP are modulated by motion, disparity and vergence eye-posture. In addition, we have shown that these variables are encoded independently, but allow recovering egocentric distance at a population level. This encoding scheme is reminiscent of the gain-field neurons for conjunctive eye-position^82^ and likely includes speed tuning as well. This mechanism could provide an answer to how various types of visual constancies, such as speed, depth and size constancy^83–85^ are implemented at a neural level.

## Methods

### Animals

Two adult rhesus macaques (Macaca mulatta: 9.2kg and 9.5kg) have been used for this study. All treatments regarding animal care were in accordance with German and international published guidelines on the use of animals in research (European Communities Council Directive 86/609/ECC).

Procedures, materials and methods are described in more detail in Bremmer et al. (2013)^33^. In short, animals were implanted with a head-holding device, binocular scleral search coils and a recording chamber under general anesthesia. Chambers were located over the Parietal sulcus based on MRI scans for each animal. In one animal the recording chamber was placed over the left cortical hemisphere, in the other it was placed over the right cortical hemisphere.

### Stimuli

We used a CRT video projector (Electrohome ECP 4100, Electrohome, Canada) to back-project the visual stimuli onto a screen that subtended the central 70×70 deg of the animal’s visual field at 48 cm. The stimuli, random dot pattern, were presented stereoscopically via LCD shutter goggles (Silicon Graphics, customized, US) at a rate of 120 Hz interlaced, meaning that each eye was shown the stimuli at 60 Hz. Each trial started with the presentation of a central fixation target (800 ms) at one of three egocentric distances corresponding to a vergence angle of 5.5°, 3.5° and 1.5° relative to the animal or -2°, 0° and +2° relative to the screen. These values correspond geometrically to distances of 31 cm, 48 cm and 101 cm relative to the animal, i.e., within, at the limit and beyond peripersonal space of a macaque monkey^86^. After the fixation period, a stationary Random Dot Pattern (RDP) was presented for 250 ms at 1 of 7 possible distances in pseudorandom order, defined by a disparity ranging from -3° to +3° relative to the display screen. These disparities corresponded geometrically to distances of 27, 31, 38, 48, 65, 101 and 229 cm relative to the animal (Figure 1). As for vergence, these values were within, at the limit of and beyond peripersonal space. The RDP then started moving along a circular path in the fronto-parallel plane^87^ for 2000 ms (1 cycle: 1600 ms). Stimuli had a constant retinal speed and were corrected for the inter-ocular delay introduced by the sequential stereoscopic system.

### Procedure

During experiments, monkeys were seated comfortably in a primate chair with their heads fixed. For each recording session, a tungsten electrode (impedance 1-2 MΩ at 1 KHz) was inserted into the lateral bank of the IntraParietal Sulcus (IPS). The electrode was lowered using a hydraulic microdrive (Narishige, Tokyo, Japan) until a neuron could be isolated over background activity. Area VIP was identified by the depth and position of the electrode within the IPS as well as physiological properties of the area (direction selectivity to the visual stimulus). We recorded a total of 113 neurons across sessions (89 from monkey C and 24 from monkey H).

The combination of 3 fixation distances (vergence) and 7 stimulus distances (disparity) corresponded to 21 stimulus conditions that were presented in pseudorandomized order across trials. Monkeys were required to maintain fixation of the central target throughout a given trial (3050 ms), at which point they received a liquid reward. We only analyzed trials where the animals successfully maintained fixation within 1 deg of visual angle during the whole course of the trial. We recorded approximately 9±2 successful trials per condition and neuron.

### Analyses

Raw spike count rate was used to quantify neural activity in the different periods of a trial. The temporal interval ranging from -300 to 0 ms relative to the onset of the stationary RDP was considered as *fixation period*. Here, only the fixation target was visible on the screen. The interval ranging from 50 to 250 ms after stationary stimulus onset was considered as *stationary period*. Importantly, this period already contained disparity information of the stimulus. For the analysis of the motion induced response we discarded the first quarter cycle of the circular pathway movement of the stimulus (400 ms) to avoid transient activity related to movement onset. Hence, we defined the temporal interval from 400ms to 2000ms after movement onset (i.e. one complete cycle) as the *motion period*.

#### Directional selectivity

Motion direction selectivity was assessed separately for each combination of vergence and stimulus disparity (21 conditions). For each condition, all the corresponding trials were stacked and binned in 50 ms width (corresponding to 22.5 degrees). Then by implementing a Rayleigh test, we checked if the cell has a directional preference (p<0.05) To assess the stability of directional tuning across various conditions, we computed the pairwise difference in preferred directions for each cell. We then compared this distribution with two other distributions. First, with a similar distribution computed on a randomly shuffled dataset. This distribution indicates differences expected if preferred directions were unstable across conditions. Second, we resampled the dataset *within* conditions. Here differences in preferred directions occur because of the noisiness of the neurons responses and indicates expected differences if their preferred directions remained perfectly stable across conditions.

#### Disparity sign selectivity

We used the Kruskal-Wallis test to measure the disparity selectivity of each neuron. Since a majority of our cells exhibited a preference for either near or far disparity, we further quantified their disparity sign selectivity using a Depth Sign Discriminant Index (DSDI) ^47,48^:

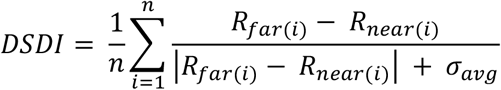

With *R*_*far*(*i*)_ and *R*_near(*i*)_ the cells firing rate in pairs of disparities of equal value but opposite sign (e.g., ±1 deg); and *σ*_*avg*_ the average standard deviation of the two responses. The DSDI quantifies how well a neuron is tuned to far or near disparities relative to the ongoing activity of the cell; it ranges from -1 (strong preference for near disparity) to +1 (strong preference for far disparity). To determine if the observed index is significantly different from randomness, we performed a permutation test by randomly shuffling the disparity sign 1 million times. We considered an index value significant if it was outside of the 95% confidence interval of the random permutated distribution. Since the tested disparities were identical in space, but not around each fixation distance in our experiment, for computing a general index we used trials when the monkey was fixating at the screen distance, in which we have 3 symmetrical disparities around the fixation point (±1°, ±2° and ±3°).

#### Vergence selectivity

First we performed Kruskal-Wallis tests for each stimulus period separately to assess whether a cell’s activity was significantly modulated by vergence angles. To identify the vergence preference (near or far) for each cell, we computed a Vergence Distance Discriminant Index (VDDI), by analogy with the DSDI:

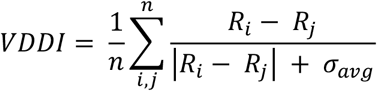

With *i,j* all pairs of different vergence angles and σ_*avg*_ is the average standard deviation of the two responses.

#### Gaussian Mixture Models

To test whether the distributions of DSDI and VDDI were better explained by a unimodal or bimodal distribution, we used a Leave-One-Out cross-validation technique: we fitted 2 Gaussian Mixture Models including respectively 1 or 2 components to all but 1 datapoint. We then computed the likelihood ratio of the unfitted datapoint according to the 2 models. We iterated this process for every point of the dataset and summed up the log likelihood ratios to compare the overall performance of one model as compared to the other.

#### Distance tuning

If neurons were tuned to distance, we anticipated unchanged activity despite variation in vergence and disparity when a stimulus was presented at a constant distance. To evaluate this, we analyzed neural responses during stimulus presentation at a fixed distance, accounting for diverse combinations of vergence and disparity. This analysis was carried out individually for each of the 7 distances using a Kruskal-Wallis test. If there was no significant change in activity for any of the distances, we further checked the possibility of being significantly tuned to one specific distance (Kruskal-Wallis test, p<0.05).

### LNP model

#### Framework

To quantify the dependence of neurons’ spiking rates on a combination of variables, we fitted a Linear-Nonlinear Poisson model (LNP) to each neuron. LNP models assume that the firing rate of a neuron is a linear combination of independent variables passed through an exponential static nonlinearity. Similar models have been used to model neural activity in multiple areas including of the Posterior Parietal Cortex^42,43,49,88^. Formally, firing rate is expressed as:

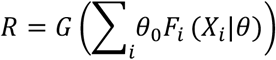

Where *i* refers to one of our 3 variables (motion direction, disparity or vergence) and the baseline, *F*_*i*_ is the neuron tuning function for each variable *i, X*_*i*_ is the value of that variable and *θ* is a set of parameters for F, G is an exponential non-linearity and *θ*_*0*_ a gain modulation factor. This model required assuming a tuning function for the 3 variables. We assumed that motion tuning was the sum of 2 von Mises functions with peaks 180° apart. Disparity was assumed to follow a Gabor function, the product of a Gaussian envelope and a sine carrier. Similarly, to previous studies^89–91^, we found that leaving the frequency component of the carrier as a free parameter led to strong overfitting. However, since VIP cells exhibit a broad tuning at low disparity frequencies always lower than 0.2cyc.deg^-1 34^, we limited that parameter to the range [0-0.2]. Finally, vergence was assumed to be a second-order polynomial, the simplest curve we could fit to our 3 vergence angles.

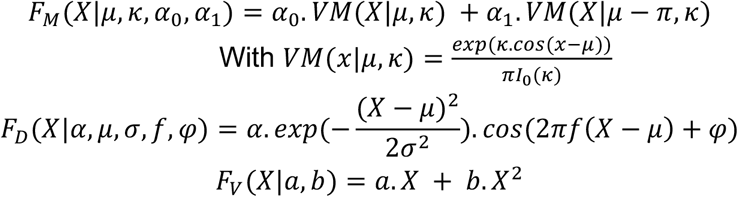

#### Estimation of the non-linearity

First we tested how the neuron’s output varied when multiple variables covary. We binned data in 16 possible motion directions and computed the mean firing rate for each of these bins (16 directions x 7 disparities x 3 vergence). Then for a given pair of variables A and B, we measured the differences in firing rate when only one variable changes: Δ_1_ = A_2_B_1_ - A_1_B_1_ and Δ_2_ = A_1_B_2_ - A_1_B_1_. If the variables contribute additively to the neuron’s activity, the difference of activity when both variables change should simply be the sum of the change when each variable changes alone: Δ_3_ = A_2_B_2_ - A_1_B_1_ = Δ_1_ + Δ_2_. Instead, we found that this latter difference was consistently higher than predicted by linear summation, and that the following equality held approximately true: log(A_2_B_2_) μ log(A_2_B_1_)+log(A_1_B_2_), indicative of an exponential non-linearity.

#### Optimization

Assuming that the firing rate is approximately a Poisson process, we treat each 1 ms time bin as independent. Therefore, the probability of a spike in each bin is approximately *r*. Δ_*t*_ and the probability of no spikes is 1 −] *r*. Δ_*t*_ (assuming that the probability of having 2 spikes within a 1 ms bin is negligible). To fit the model, we computed and summed the log-likelihood of having observed a spike, or no spike, in each time bin jointly across the entire dataset (on average 235,000 data-points). We used BADS^92^ to find the set of parameters maximizing this log-likelihood.

#### Selection

For model selection, we used a nested forward-search approach combined with 10-fold cross-validation. For each cell we divided the dataset in 10 random subsets and fitted the model to all but 1 of the subsets. We then computed the likelihood of the unfitted subset. We first fitted a model including the baseline firing rate only. We then added each variable one at a time and preserved variables only when it improved the likelihood of the model, in order of improvement of the likelihood. This allowed us to estimate jointly whether a variable contributes the cell firing-rate, and the tuning function of the cell.

#### Comparison with actual tuning curves

To estimate whether our model captured the cell’s actual pattern of activity, we performed the same analyses on predicted firing rate of the model as on the real data. Since we assume a Poisson process for generating spikes, we focus on the mean firing rate rather than the variability for comparing metrics predicted by the model and computed the DSDI and VDDI using the standard measurement of contrast: 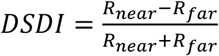 (and similarly for the VDDI). The correlation coefficient between actual DSDI and DSDI predicted by the model was 0.99 and the correlation for VDDI was 0.94.

#### Decoding

Decoding was performed using Maximum Likelihood Estimation. The encoding model provides the probability distribution *p*(*r|d,v,s*): the conditional probability of observing spike count rate *r* given the disparity *d*, vergence *v* and direction *s*. For decoding we need to reverse this problem to find the probability of each parameter given the spike count rate *p*(*d,v,s|r*):. We assumed a flat prior over stimulus values, in which case decoding can be performed by simply summing the log-probability of each stimulus value across the population:

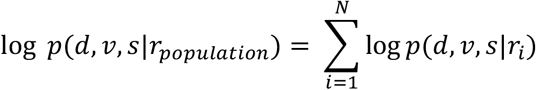

The variables associated with the maximum a posteriori will be assigned as the decoded value.

## Acknowledgment

Funded by the Deutsche Forschungsgemeinschaft (DFG, German Research Foundation) – 524696675 to B.C., the IRTG-1901 and CRC/TRR-135 (project number 222641018) to F.B, by the state ministry for research (HMWK) – Clusterproject The Adaptive Mind – TAM and EU-PLACES.

## References

1. Vallar, G., and Coslett, H.B. (2018). The parietal lobe (Academic Press).

2. Whitlock, J.R. (2017). Posterior parietal cortex. Curr. Biol. 27, R691–R695.

3. Holmes, G., and Horrax, G. (1919). Disturbances of spatial orientation and visual attention, with loss of stereoscopic vision. Arch. Neurol. Psychiatry 1, 385–407.

4. Brain, W.R. (1941). Visual orientation with special reference to lesions of the right cerebral hemisphere. Brain J. Neurol.

5. Cogan, D.G. (1965). Ophthalmic manifestations of bilateral non-occipital cerebral lesions. Br. J. Ophthalmol. 49, 281.

6. Murphy, A.P., Leopold, D.A., Humphreys, G.W., and Welchman, A.E. (2016). Lesions to right posterior parietal cortex impair visual depth perception from disparity but not motion cues. Philos. Trans. R. Soc. B Biol. Sci. 371, 20150263. 10.1098/rstb.2015.0263.

7. Duhamel, J.-R., Bremmer, F., Ben Hamed, S., and Graf, W. (1997). Spatial invariance of visual receptive fields in parietal cortex neurons. Nature 389, 845–848.

8. Schlack, A., Sterbing-D’Angelo, S.J., Hartung, K., Hoffmann, K.-P., and Bremmer, F. (2005). Multi-sensory space representations in the macaque ventral intraparietal area. J. Neurosci. 25, 4616–4625.

9. Avillac, M., Deneve, S., Olivier, E., Pouget, A., and Duhamel, J.-R. (2005). Reference frames for representing visual and tactile locations in parietal cortex. Nat. Neurosci. 8, 941–949.

10. Zipser, D., and Andersen, R.A. (1988). A back-propagation programmed network that simulates response properties of a subset of posterior parietal neurons. Nature 331, 679–684.

11. Morris, A.P., Kubischik, M., Hoffmann, K.-P., Krekelberg, B., and Bremmer, F. (2012). Dynamics of eye-position signals in the dorsal visual system. Curr. Biol. 22, 173–179.

12. Morris, A.P., and Krekelberg, B. (2019). A stable visual world in primate primary visual cortex. Curr. Biol. 29, 1471–1480.

13. Bremmer, Pouget, and Hoffmann, K.-P. (1998). Eye position encoding in the macaque posterior parietal cortex. Eur. J. Neurosci. 10, 153–160. 10.1046/j.1460-9568.1998.00010.x.

14. Wheatstone, C. (1838). XVIII. Contributions to the physiology of vision.—Part the first. On some remarkable, and hitherto unobserved, phenomena of binocular vision. Philos. Trans. R. Soc. Lond., 371–394.

15. DeAngelis, G.C. (2000). Seeing in three dimensions: the neurophysiology of stereopsis. Trends Cogn. Sci. 4, 80–90. 10.1016/S1364-6613(99)01443-6.

16. Gonzalez, F., and Perez, R. (1998). Neural mechanisms underlying stereoscopic vision. Prog. Neurobiol. 55, 191–224.

17. Howard, I.P. (2002). Seeing in depth, Vol. 1: Basic mechanisms. (University of Toronto Press).

18. Parker, A.J., and Cumming, B.G. (2001). Cortical mechanisms of binocular stereoscopic vision. Prog. Brain Res. 134, 205–216.

19. Tresilian, J.R., Mon-Williams, M., and Kelly, B.M. (1999). Increasing confidence in vergence as a cue to distance. Proc. R. Soc. Lond. B Biol. Sci. 266, 39–44.

20. Viguier, A., Clement, G., and Trotter, Y. (2001). Distance perception within near visual space. Perception 30, 115–124.

21. Foley, J.M. (1985). Binocular distance perception: egocentric distance tasks. J. Exp. Psychol. Hum. Percept. Perform. 11, 133.

22. Morrison, J.D., and Whiteside, T.C. (1984). Binocular cues in the perception of distance of a point source of light. Perception 13, 555–566.

23. Pouget, A., and Sejnowski, T.J. (1994). A neural model of the cortical representation of egocentric distance. Cereb. CORTEX-N. Y.-Oxf. Univ. Press. 4, 314–314.

24. Trotter, Y., Celebrini, S., Stricanne, B., Thorpe, S., and Imbert, M. (1992). Modulation of Neural Stereoscopic Processing in Primate Area V1 by the Viewing Distance. Science 257, 1279–1281. 10.1126/science.1519066.

25. Trotter, Y., Celebrini, S., Stricanne, B., Thorpe, S., and Imbert, M. (1996). Neural processing of stereopsis as a function of viewing distance in primate visual cortical area V1. J. Neurophysiol. 76, 2872–2885. 10.1152/jn.1996.76.5.2872.

26. Gonzalez, F., and Perez, R. (1998). Modulation of cell responses to horizontal disparities by ocular vergence in the visual cortex of the awake macaca mulatta monkey. Neurosci. Lett. 245, 101–104. 10.1016/S0304-3940(98)00191-8.

27. Cumming, B.G., and Parker, A.J. (1999). Binocular Neurons in V1 of Awake Monkeys Are Selective for Absolute, Not Relative, Disparity. J. Neurosci. 19, 5602–5618. 10.1523/JNEURO-SCI.19-13-05602.1999.

28. Cumming, B.G., and DeAngelis, G.C. (2001). The physiology of stereopsis. Annu. Rev. Neurosci. 24, 203–238.

29. Minini, L., Parker, A.J., and Bridge, H. (2010). Neural Modulation by Binocular Disparity Greatest in Human Dorsal Visual Stream. J. Neurophysiol. 104, 169–178. 10.1152/jn.00790.2009.

30. Neri, P., Bridge, H., and Heeger, D.J. (2004). Stereoscopic processing of absolute and relative disparity in human visual cortex. J. Neurophysiol. 92, 1880–1891.

31. Gnadt, J.W., and Mays, L.E. (1995). Neurons in monkey parietal area LIP are tuned for eye-movement parameters in three-dimensional space. J. Neurophysiol. 73, 280–297. 10.1152/jn.1995.73.1.280.

32. Genovesio, A., and Ferraina, S. (2004). Integration of retinal disparity and fixation-distance related signals toward an egocentric coding of distance in the posterior parietal cortex of primates. J. Neurophysiol. 91, 2670–2684.

33. Bremmer, F., Schlack, A., Kaminiarz, A., and Hoffmann, K.-P. (2013). Encoding of movement in near extrapersonal space in primate area VIP. Front. Behav. Neurosci. 7, 8.

34. Yang, Y., Liu, S., Chowdhury, S.A., DeAngelis, G.C., and Angelaki, D.E. (2011). Binocular disparity tuning and visual–vestibular congruency of multisensory neurons in macaque parietal cortex. J. Neurosci. 31, 17905–17916.

35. Bhattacharyya, R., Musallam, S., and Andersen, R.A. (2009). Parietal reach region encodes reach depth using retinal disparity and vergence angle signals. J. Neurophysiol. 102, 805–816.

36. Sakata, H., Taira, M., Kusunoki, M., Murata, A., Tanaka, Y., and Tsutsui, K. (1998). Neural coding of 3D features of objects for hand action in the parietal cortex of the monkey. Philos. Trans. R. Soc. B Biol. Sci. 353, 1363–1373.

37. Sakata, H., Taira, M., Kusunoki, M., Murata, A., Tsutsui, K., Tanaka, Y., Shein, W.N., and Miyashita, Y. (1999). Neural representation of three-dimensional features of manipulation objects with stereopsis. Exp. Brain Res. 128, 160–169.

38. Filippini, M., Breveglieri, R., Hadjidimitrakis, K., Bosco, A., and Fattori, P. (2018). Prediction of reach goals in depth and direction from the parietal cortex. Cell Rep. 23, 725–732.

39. Bremmer, F., Schlack, A., Duhamel, J.-R., Graf, W., and Fink, G.R. (2001). Space coding in primate posterior parietal cortex. Neuroimage 14, S46–S51.

40. Andersen, R.A. (1997). Multimodal integration for the representation of space in the posterior parietal cortex. Philos. Trans. R. Soc. Lond. B. Biol. Sci. 352, 1421–1428.

41. Sakata, H., Shibutani, H., and Kawano, K. (1980). Spatial properties of visual fixation neurons in posterior parietal association cortex of the monkey. J. Neurophysiol. 43, 1654–1672. 10.1152/jn.1980.43.6.1654.

42. Park, I.M., Meister, M.L., Huk, A.C., and Pillow, J.W. (2014). Encoding and decoding in parietal cortex during sensorimotor decision-making. Nat. Neurosci. 17, 1395–1403.

43. Hardcastle, K., Maheswaranathan, N., Ganguli, S., and Giocomo, L.M. (2017). A multiplexed, heterogeneous, and adaptive code for navigation in medial entorhinal cortex. Neuron 94, 375–387.

44. Bremmer, F., Klam, F., Duhamel, J.-R., Ben Hamed, S., and Graf, W. (2002). Visual–vestibular interactive responses in the macaque ventral intraparietal area (VIP). Eur. J. Neurosci. 16, 1569–1586.

45. Colby, C.L., Duhamel, J.-R., and Goldberg, M.E. (1993). Ventral intraparietal area of the macaque: anatomic location and visual response properties. J. Neurophysiol. 69, 902–914.

46. Roy, J., Komatsu, H., and Wurtz, R. (1992). Disparity sensitivity of neurons in monkey extrastriate area MST. J. Neurosci. 12, 2478–2492. 10.1523/JNEUROSCI.12-07-02478.1992.

47. Nadler, J.W., Angelaki, D.E., and DeAngelis, G.C. (2008). A neural representation of depth from motion parallax in macaque visual cortex. Nature 452, 642–645.

48. Nadler, J.W., Nawrot, M., Angelaki, D.E., and DeAngelis, G.C. (2009). MT neurons combine visual motion with a smooth eye movement signal to code depth-sign from motion parallax. Neuron 63, 523–532.

49. Lakshminarasimhan, K.J., Avila, E., Pitkow, X., and Angelaki, D.E. (2023). Dynamical latent state computation in the male macaque posterior parietal cortex. Nat. Commun. 14, 1832.

50. Cooke, D.F., Taylor, C.S.R., Moore, T., and Graziano, M.S.A. (2003). Complex movements evoked by microstimulation of the ventral intraparietal area. Proc. Natl. Acad. Sci. 100, 6163–6168. 10.1073/pnas.1031751100.

51. Graziano, M.S.A., and Cooke, D.F. (2006). Parieto-frontal interactions, personal space, and defensive behavior. Neuropsychologia 44, 2621–2635. 10.1016/j.neuropsychologia.2005.09.011.

52. de Borst, A.W., and de Gelder, B. (2022). Threat Detection in Nearby Space Mobilizes Human Ventral Premotor Cortex, Intraparietal Sulcus, and Amygdala. Brain Sci. 12, 391. 10.3390/brainsci12030391.

53. Bremmer, F., Schlack, A., Shah, N.J., Zafiris, O., Kubischik, M., Hoffmann, K.-P., Zilles, K., and Fink, G.R. (2001). Polymodal motion processing in posterior parietal and premotor cortex: a human fMRI study strongly implies equivalencies between humans and monkeys. Neuron 29, 287–296.

54. Foster, C., Sheng, W.-A., Heed, T., and Ben Hamed, S. (2022). The macaque ventral intraparietal area has expanded into three homologue human parietal areas. Prog. Neurobiol. 209, 102185. 10.1016/j.pneurobio.2021.102185.

55. Roy, J.-P., and Wurtz, R.H. (1990). The role of disparity-sensitive cortical neurons in signalling the direction of self-motion. Nature 348, 160–162.

56. Sprague, W.W., Cooper, E.A., Tošić, I., and Banks, M.S. (2015). Stereopsis is adaptive for the natural environment. Sci. Adv. 1, e1400254. 10.1126/sciadv.1400254.

57. Ben Hamed, S., Duhamel, J.-R., Bremmer, F., and Graf, W. (2001). Representation of the visual field in the lateral intraparietal area of macaque monkeys: a quantitative receptive field analysis. Exp. Brain Res. 140, 127–144. 10.1007/s002210100785.

58. Morris, A.P., Bremmer, F., and Krekelberg, B. (2013). Eye-position signals in the dorsal visual system are accurate and precise on short timescales. J. Neurosci. 33, 12395–12406.

59. Bremmer, F., Distler, C., and Hoffmann, K.-P. (1997). Eye Position Effects in Monkey Cortex. II. Pursuit- and Fixation-Related Activity in Posterior Parietal Areas LIP and 7A. J. Neurophysiol. 77, 962–977. 10.1152/jn.1997.77.2.962.

60. Bremmer, F., Ilg, U.J., Thiele, A., Distler, C., and Hoffmann, K.-P. (1997). Eye Position Effects in Monkey Cortex. I. Visual and Pursuit-Related Activity in Extrastriate Areas MT and MST. J. Neurophysiol. 77, 944–961. 10.1152/jn.1997.77.2.944.

61. Bremmer, F., Graf, W., Hamed, S.B., and Duhamel, J.-R. (1999). Eye position encoding in the macaque ventral intraparietal area (VIP). Neuroreport 10, 873–878.

62. Xu, Y., Wang, X., Peck, C., and Goldberg, M.E. (2011). The time course of the tonic oculomotor proprioceptive signal in area 3a of somatosensory cortex. J. Neurophysiol. 106, 71–77. 10.1152/jn.00668.2010.

63. Ogle, K.N. (1950). Researches in binocular vision. (WB Saunders).

64. Mayhew, J.E.W., and Longuet-Higgins, H.C. (1982). A computational model of binocular depth perception. Nature 297, 376–378.

65. Bishop, P.O. (1989). Vertical disparity, egocentric distance and stereoscopic depth constancy: a new interpretation. Proc. R. Soc. Lond. B Biol. Sci. 10.1098/rspb.1989.0059.

66. Backus, B.T., Banks, M.S., van Ee, R., and Crowell, J.A. (1999). Horizontal and vertical disparity, eye position, and stereoscopic slant perception. Vision Res. 39, 1143–1170. 10.1016/S0042-6989(98)00139-4.

67. Cumming, B.G., Johnston, E.B., and Parker, A.J. (1991). Vertical disparities and perception of three-dimensional shape. Nature 349, 411–413. 10.1038/349411a0.

68. Vienne, C., Plantier, J., Neveu, P., and Priot, A.-E. (2016). The Role of Vertical Disparity in Distance and Depth Perception as Revealed by Different Stereo-Camera Configurations. -Percept. 7, 204166951668130. 10.1177/2041669516681308.

69. Linton, P. (2020). Does vision extract absolute distance from vergence? Atten. Percept. Psychophys. 82, 3176–3195. 10.3758/s13414-020-02006-1.

70. Chopin, A., Levi, D., Knill, D., and Bavelier, D. (2016). The absolute disparity anomaly and the mechanism of relative disparities. J. Vis. 16, 2–2.

71. Blakemore, C. (1970). The range and scope of binocular depth discrimination in man. J. Physiol. 211, 599–622.

72. Backus, B. (2001). Perceptual metamers in stereoscopic vision. Adv. Neural Inf. Process. Syst. 14.

73. Foley, J.M. (1980). Binocular distance perception. Psychol. Rev. 87, 411.

74. Wheatstone, C. (1852). I. The Bakerian Lecture.—Contributions to the physiology of vision.—Part the second. On some remarkable, and hitherto unobserved, phenomena of binocular vision (continued). Philos. Trans. R. Soc. Lond., 1–17.

75. Caziot, B., Valsecchi, M., Gegenfurtner, K.R., and Backus, B.T. (2015). Fast perception of binocular disparity. J. Exp. Psychol. Hum. Percept. Perform. 41, 909.

76. Caziot, B., Backus, B.T., and Lin, E. (2017). Early dynamics of stereoscopic surface slant perception. J. Vis. 17, 4–4.

77. Caziot, B., Rolfs, M., and Backus, B.T. (2023). Orienting attention across binocular disparity. PNAS Nexus 2, pgad314.

78. Finlayson, N.J., and Grove, P.M. (2015). Visual search is influenced by 3D spatial layout. Atten. Percept. Psychophys. 77, 2322–2330. 10.3758/s13414-015-0924-3.

79. Caziot, B., and Backus, B.T. (2015). Stereoscopic offset makes objects easier to recognize. PloS One 10, e0129101.

80. Burr, D.C., and Morrone, M.C. (2011). Spatiotopic coding and remapping in humans. Philos. Trans. R. Soc. B Biol. Sci. 366, 504–515. 10.1098/rstb.2010.0244.

81. Maunsell, J.H.R., and Cook, E.P. (2002). The role of attention in visual processing. Philos. Trans. R. Soc. Lond. B. Biol. Sci. 10.1098/rstb.2002.1107.

82. Andersen, R.A., Essick, G.K., and Siegel, R.M. (1985). Encoding of Spatial Location by Posterior Parietal Neurons. Science 230, 456–458. 10.1126/science.4048942.

83. McKee, S.P., and Smallman, H.S. (1998). Size and speed constancy.

84. Collett, T.S., and Parker, A.J. (1998). 15 Depth constancy. Percept. Constancy Why Things Look They Do, 409.

85. Allison, R.S., and Wilcox, L.M. (2021). Stereoscopic depth constancy from a different direction. Vision Res. 178, 70–78. 10.1016/j.visres.2020.10.003.

86. Caggiano, V., Fogassi, L., Rizzolatti, G., Thier, P., and Casile, A. (2009). Mirror Neurons Differentially Encode the Peripersonal and Extrapersonal Space of Monkeys. Science 324, 403–406. 10.1126/science.1166818.

87. Bremmer, F., Kubischik, M., Pekel, M., Hoffmann, K.-P., and Lappe, M. (2010). Visual selectivity for heading in monkey area MST. Exp. Brain Res. 200, 51–60. 10.1007/s00221-009-1990-3.

88. Mao, D., Avila, E., Caziot, B., Laurens, J., Dickman, J.D., and Angelaki, D.E. (2021). Spatial modulation of hippocampal activity in freely moving macaques. Neuron 109, 3521–3534.

89. DeAngelis, G.C., and Uka, T. (2003). Coding of Horizontal Disparity and Velocity by MT Neurons in the Alert Macaque. J. Neurophysiol. 89, 1094–1111. 10.1152/jn.00717.2002.

90. Prince, S.J.D., Cumming, B.G., and Parker, A.J. (2002). Range and Mechanism of Encoding of Horizontal Disparity in Macaque V1. J. Neurophysiol. 87, 209–221. 10.1152/jn.00466.2000.

91. Prince, S.J.D., Pointon, A.D., Cumming, B.G., and Parker, A.J. (2002). Quantitative Analysis of the Responses of V1 Neurons to Horizontal Disparity in Dynamic Random-Dot Stereograms. J. Neurophysiol. 87, 191–208. 10.1152/jn.00465.2000.

92. Acerbi, L., and Ma, W.J. (2017). Practical Bayesian optimization for model fitting with Bayesian adaptive direct search. Adv. Neural Inf. Process. Syst. 30.

93. Busettini, C., Miles, F.A., and Krauzlis, R.J. (1996). Short-latency disparity vergence responses and their dependence on a prior saccadic eye movement. J. Neurophysiol. 75, 1392–1410. 10.1152/jn.1996.75.4.1392.

