## Supplementary for "Coding of egocentric distance in the macaque ventral intraparietal area"

**Supplementary Material**

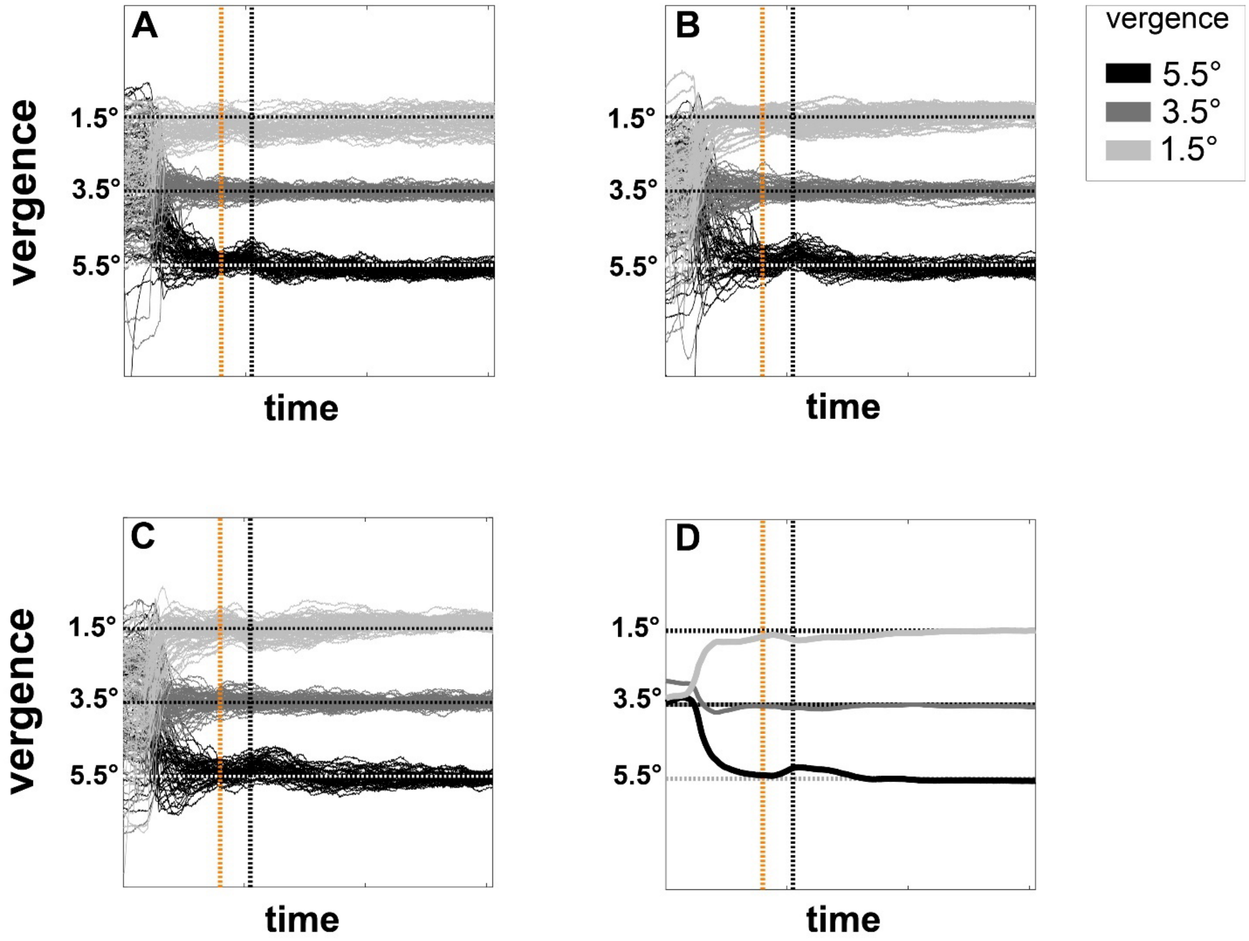


**Figure S1: A-C:** Vergence eye posture in the course of three representative sessions. Different colors representing different vergence angle. **D:** Average vergence angle recorded across sessions. The standard error is too small to be visible in this figure. Dotted horizontal lines illustrate the vergence demand of the fixation dot. Two vertical lines showing the start of stationary (orange) and motion period (black).

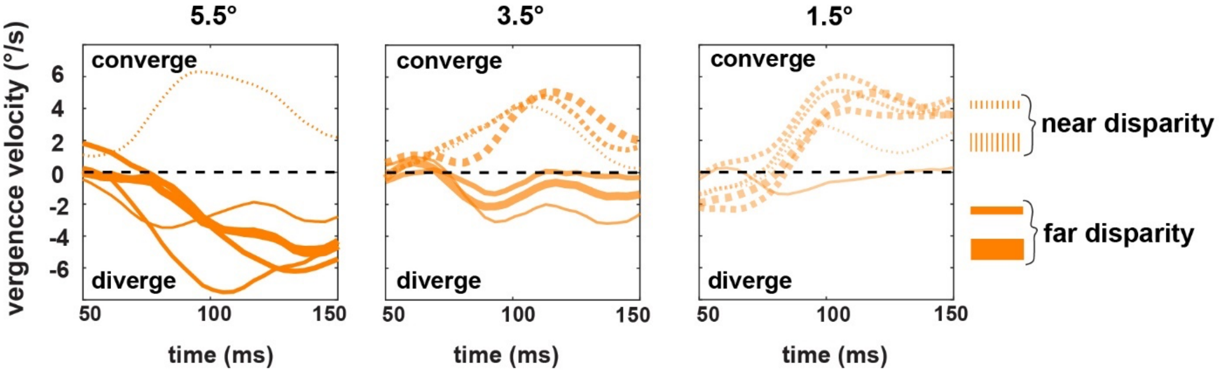


**Figure S2:** Mean vergence velocity from 50 to 150 ms after stimulus onset (during the stationary period) as a function of disparity. Each panel represent data from one vergence angle (presented at the top). Dotted lines depict crossed disparities and solid lines uncrossed disparities, and thicker lines represent higher disparity values. Positive and negative values on the y-axis denote convergence and divergence eye movement, respectively.

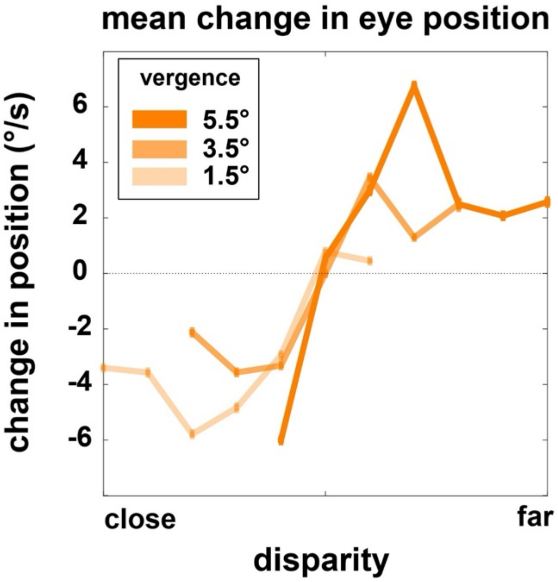


**Figure S3**: Average change in eye position between 76 and 110 ms after stimulus onset as a function of disparity and vergence angle.

Appearance of the static RDP induced a small reflex-like vergence response. This response explains the small deviation in the vergence angle that appears roughly 70 ms after presentation of the stationary stimulus (see Figure S1). Since disparities were inversely related to vergence angle, the resulting deviations in the vergence postures converge towards the mean stimulus disparity (see Figure S2), but return to fixation vergence demand briefly after motion onset. This response exhibits the expected non-linear tuning between disparity and vergence response demonstrative of the open-loop nature of this response[93] (see Figure S3). The first quarter of the motion period was discarded from analyses.

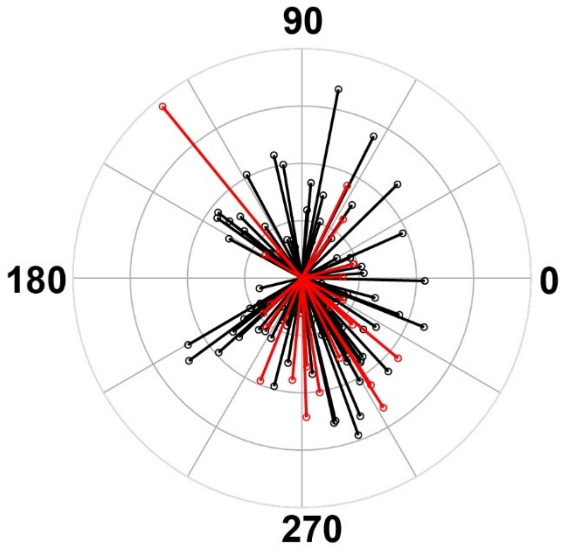


**Figure S4:** Preferred direction of each individual cell. Each color represents data from one monkey.

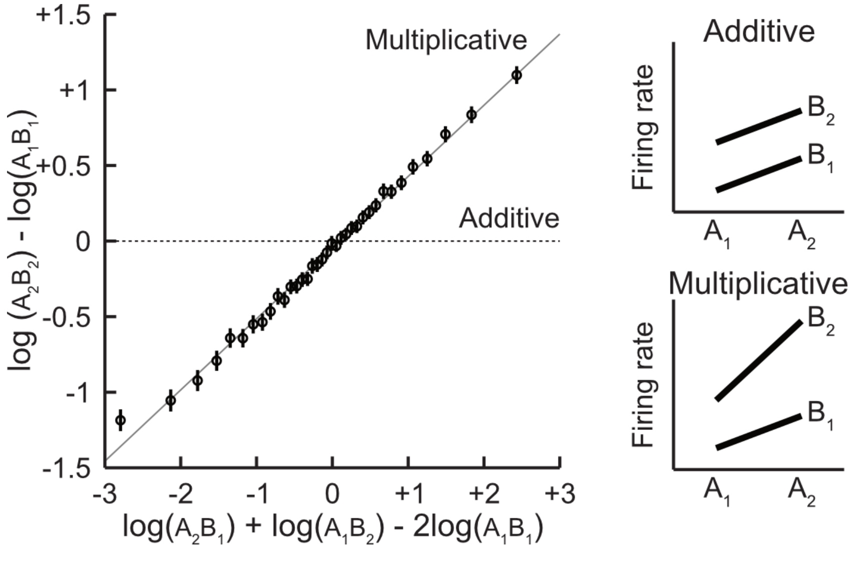


**Figure S5:** Evidence for a multiplicative encoding of task variables. Here changes in firing rates activity were computed between all pairs of matching conditions within neurons.

We tested whether variables interact linearly or multiplicatively by computing the firing rate within bins where 2 variables are jointly changing. Specifically, if variables A and B are changing between conditions $A_1$ to $A_2$ and $B_1$ to $B_2$, then if variables A and B are contributing additively to the activity of the neuron the change in firing rate between conditions $A_1B_1$ and $A_2B_2$ should be equal to the sum of the change when either variable changed alone, or $A_2B_1$ - $A_1B_1$ and $A_1B_2$ - $A_1B_1$. Instead, if variables contribute multiplicatively to the neuron's activity, we expect to observe a linear relationship between $\log(A_2B_2) - \log(A_1B_1)$ and $\log(A_2B_1) + \log(A_1B_2) - 2\log(A_1B_1)$. Figure S5 plots the average of this measure across all possible pairs of matching conditions and neurons, and show that this relationship holds approximately true, demonstrating a multiplicative interaction between variables.

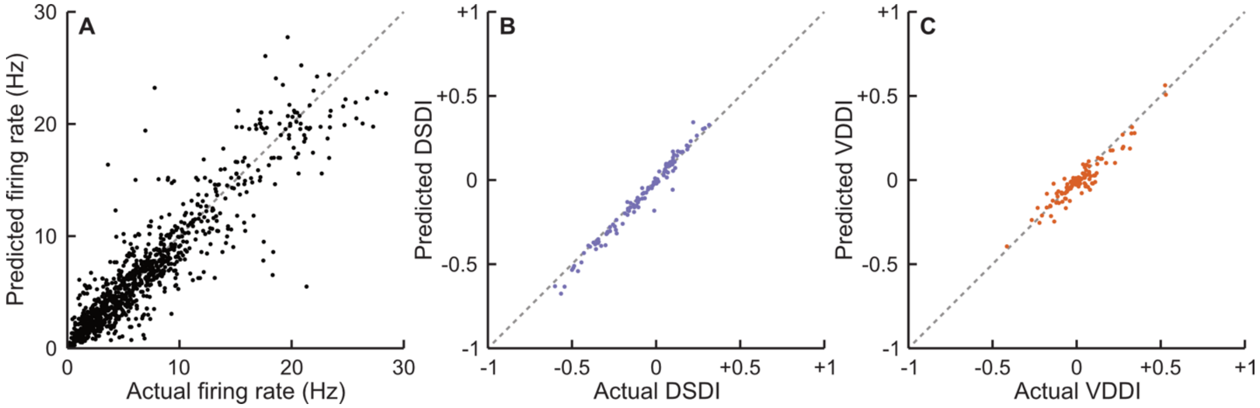


**Figure S6**: **A:** Firing rate predicted by the LNP model as a function of actual firing rate across all neurons and conditions. **B:** DSDI predicted by the model as a function of actual DSDI. **C:** Same for VDDI.

Firing rates and indices DSDI and VDDI predicted by the model fits were highly correlated with actual measurements. Note that here DSDI and VDDI were computed using the standard contrast measurement instead of a metric that includes a measurement of responses variability (see Methods).
